# Acidic vacuole-containing organisms are a majority of the eukaryotic microbial community in oligotrophic Argo Basin waters (eastern Indian Ocean)

**DOI:** 10.1101/2025.08.14.670408

**Authors:** Karen E. Selph, Natalia Yingling, Claudia Traboni, Michael R. Landry

**Author notes:** Corresponding author: E-mail address (K.E. Selph).

## Abstract

The Argo Basin of the eastern Indian Ocean in austral summer (February 2022) was characterized by warm (28.5-30.6°C), oligotrophic surface waters (nitrate and phosphate ≤0.1 µM), with relatively shallow mixed layers and deep chlorophyll biomass maxima. From euphotic zone depth-resolved samples analyzed by for DNA and acid vacuole staining (Hoechst and LysoTracker Green) by ship-board flow cytometry, we found that autotrophic populations were dominated by *Prochlorococcus*, followed by mixotrophs (58 and 28% of autotrophic community biomass, respectively), with only 14% obligate phototrophic phytoplankton (i.e., plastidic cells without acid vacuole fluorescence). Acid vacuole-containing microbes (mixotrophs and heterotrophs) were 34% of the microbial community, and 80% of the eukaryotic biomass. In shallow waters, the eukaryotic chlorophyll-containing community was comprised of pico-sized obligate phototrophs and mixotrophs (233-325 cells mL^-1^), nano-sized obligate phototrophs and mixotrophs (72 and 374 cells mL^-1^, respectively), with all groups increasing several-fold in the deep chlorophyll maxima. Mixotrophs were a higher proportion of the chlorophyll-containing community in the shallow nutrient-poor mixed layer, consistent with a nutrient-acquisition argument for their prevalence. Heterotrophic eukaryotes averaged 524 ± 36 cells mL^-1^ in the euphotic zone, changing little with depth and showing a significant positive relationship with *Prochlorococcus*, but not any other group. In contrast, mixotrophs were positively correlated with heterotrophic bacteria, but not with *Prochlorococcus*. Overall, the high proportion of mixotrophs in the microbial community may channel more productivity to higher trophic levels than expected given the region’s nutrient-poor status.

## 1. Introduction

Marine microbes are comprised of autotrophs, heterotrophs, and mixotrophs. The first trophic level, the autotrophs or phytoplankton, are classically defined as organisms that only obtain their nutrition by photosynthesis (phototrophy), using solar energy to fix organic carbon and taking up dissolved inorganic nutrients. More broadly, phototrophs that require osmotrophic uptake of small organic molecules like vitamins might still be considered autotrophs. However, we now understand that many eukaryotes with light-harvesting pigments are also able to consume other microbes, making them mixotrophs – capable of obtaining nutrition by both photo- and phagotrophy (Mitra et al., 2014). These “mixoplankton” (Flynn et al., 2019) are widespread in marine systems (e.g., Unrein et al., 2007; Zubkov and Tarran, 2008; Stukel et al., 2011; Hartmann et al., 2013; Leles et al., 2018), suggesting that mixotrophy is a common nutritional strategy for life at the lowest trophic levels. Consequently, our traditional view of pelagic food webs may require re-imagining to reflect actual energy flows.

A difficult issue for determining the importance and prevalence of mixotrophs in the natural environment is that methods for quantifying autotrophic and heterotrophic eukaryotes generally distinguish them based on presence or absence of chlorophyll (chloroplasts) (e.g., Sherr et al., 1993). Mixotrophs, however, have chloroplasts or retain their endosymbiont’s plastids while also functioning as phagotrophs. One unambiguous determinant of feeding by heterotrophic eukaryotes is the presence of acidic vacuoles following phagocytosis, i.e., prey particle uptake (Mills, 2020). Vacuoles that enclose prey particles undergo intracellular cyclosis or cytoplasmic streaming. In a process that starts with neutral pH, the vacuoles acidify rapidly (∼pH 3) when fusing with acidosomes, then lysosomes, as various enzymes (mainly acid hydrolases active at pH 5.0) are introduced to break down the ingested prey (Allen and Fok, 1983). In the digestive process that lasts just a few minutes, the final step is fusion of cell membranes to expel the undigested debris (Fok et al., 1982).

The intrinsic property of pH change during phagocytosis can be exploited to identify which components of a natural mixed microbial community are functionally phagotrophic using a fluorescent dye, LysoTracker Green (LTG), that accumulates in acidic vacuoles (Rose et al., 2004). When LTG is added to a natural sample of microbes, we assume that those exhibiting an LTG signal have digestive vacuoles and function as phagotrophs, regardless of whether they contain chlorophyll in their cells (Fig. 1). This methodology is limited, however, to analysis of living cells as LTG leaks out of the cell once they die or are preserved. Flow cytometry is an established approach for rapid enumeration and fluorescence analysis of bacteria and protists in the marine environment (Vives-Rigo et al., 2000; Legendre et al., 2001; Selph, 2021) and provides the means for evaluating phagotrophic potential of natural communities by the LTG staining method.

**Figure 1.**
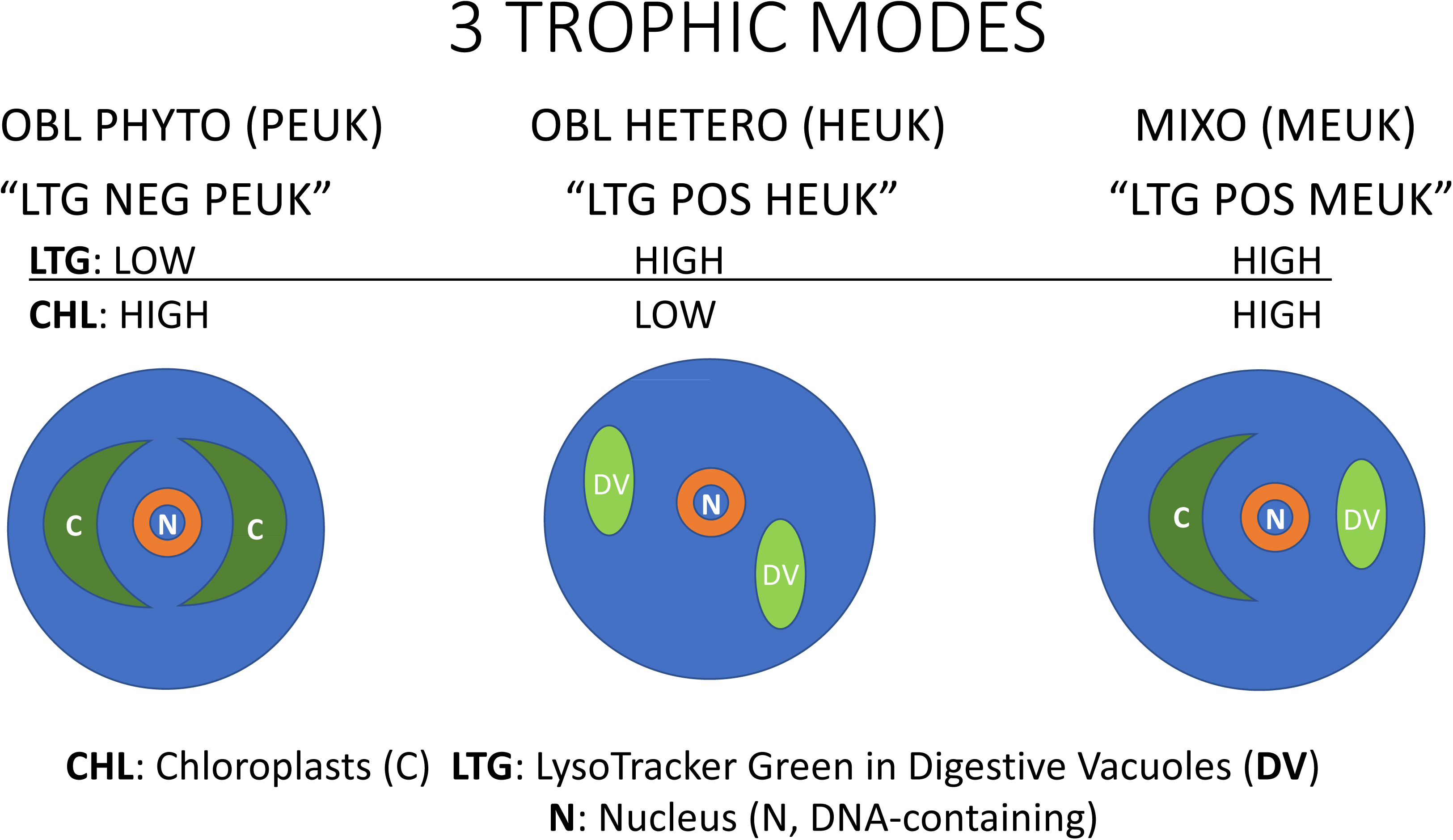
Diagram of 3 trophic modes for microbial eukaryotic plankton with reference to the magnitude of their LysoTracker Green (LTG) and Chlorophyll *a* (CHL) fluorescence. The trophic modes are obligate photosynthetic eukaryotes (PEUK), obligate heterotrophic eukaryotes (HEUK), and mixotrophic eukaryotes (MEUK). LTG negative phytoplankton are those with low LTG fluorescence (absence of digestive vacuoles) and high chlorophyll fluorescence (“LTG NEG PEUK”). LTG positive cells are divided into those with chlorophyll fluorescence, and thus presumed mixotrophs (“LTG POS MEUK”) and those with low chlorophyll fluorescence and thus presumed heterotrophs (“LTG POS HEUK”).

Some research has been done using LTG to mark acidic vacuoles of marine eukaryotes in cultures and grow-out experiments (Rose et al., 2004; Vasquez-Domingues et al., 2005; Sintes and Del Giorgio, 2010), as well as in coastal surface populations (Rose et al., 2004; Lefort and Gasol, 2014; Anderson et al., 2017). However, depth-profile data for open-ocean communities remain scarce (Sato and Hashihama, 2019). Here, we describe flow cytometric analyses using LTG to examine mixotrophy within the microbial community of oligotrophic waters overlying the Argo Abyssal Plain (hereafter, Argo Basin) off NW Australia, in the only know spawning region for Southern Bluefin Tuna. This research was conducted during the BLOOFINZ-IO (Bluefin Larvae in Oligotrophic Ocean Foodwebs, Investigations of Nutrients to Zooplankton – Indian Ocean) expedition in January-March 2022, and contributes to understanding of factors that structure food webs in the larval habitat. We found that ∼40% (abundance) of the chlorophyll-bearing eukaryotes in that region show evidence of acidic vacuole fluorescence and should be considered functional mixotrophs.

## 2. Methods

### 2.1 Study Description

The Argo Basin study area in the eastern Indian Ocean was sampled on cruise RR2201 of R/V *Roger Revelle* in February 2022 (Landry et al., this issue). Initial stations were chosen based on physical and biological attributes from site surveys and satellite imagery, geared toward finding areas that might have larval Southern Bluefin Tuna. The initial station (Day 1) involved deployment of a free-drifting in-situ array containing euphotic zone samples from that location’s CTD cast. The array had a surface float and 3x1-m holey sock drogue, and was followed to its new location 24 h later, when it was recovered, with new CTD samples at that location prepared (Day 2) and the in situ array re-deployed with those new samples. This activity was repeated for 3-5 days, following the in-situ array, creating a “Cycle” of experiments on water sampled with this Lagrangian sampling scheme (Landry et al., 2009, this issue).

At each station, water samples were collected at ∼02:00 local time at 6-8 depths in the euphotic zone from Niskin bottles mounted on a 12-place rosette. The rosette was equipped with a Sea Bird Scientific 911 CTD with sensors for temperature, conductivity, chlorophyll fluorescence (Wet Labs FLRTD-1156), and oxygen. The Wet Labs fluorometer was calibrated with discrete samples to give chlorophyll *a* (Chl*a*, µg L^-1^). A Biospherical Model QSP200L4S PAR (Photosynthetically Active Radiation) 2-pi sensor was mounted on the rosette for light measurements during daytime hours. All stations included samples for microbial community composition and nutrients, as well as samples designed to fully characterize the rest of the biota (e.g., net tows for zooplankton and larval fish abundance; Landry et al., this issue). Nutrient samples were 0.2-µm filtered directly from the rosette bottles, frozen, and analyzed at the Ocean Data Facility (Scripps Institution of Oceanography Ocean) using an autoanalyzer for nitrate+nitrite, ammonium, phosphate, and silicic acid (Becker et al., 2019). Microbial community composition was determined from a combination of flow cytometry (described herein) and epifluorescence microscopy (summarized below and described in detail in Yingling et al., this issue).

### 2.2 Microbial Community Characterization: Flow Cytometry

Within 2 h of collection, samples were prepared for shipboard analyses on a Beckman Coulter CytoFlex S flow cytometer (Selph, 2021). For determination of acidic vacuole-containing organisms, only live samples were analyzed since preserved cells do not retain the LysoTracker Green (LTG) fluorescence. Samples were stained with LTG (75 nM final concentration after Rose et al., 2004) and Hoechst 34580 (1 µg mL^-1^) for DNA staining for 10 min, then analyzed on the flow cytometer for 10 min at 50 µL min^-1^ (500 µL sample volume). Thresholding was set on fluorescence signals from Hoechst DNA or chlorophyll, so that cells with either would be counted. Fluorescent calibration beads (1 µm) were also run with each set of samples. Three prokaryote populations and three eukaryotic categories were identified: non-pigmented or heterotrophic bacteria (HBACT), *Prochlorococcus* (PRO), *Synechococcus* (SYN), heterotrophic eukaryotes (HEUK), obligate photosynthetic eukaryotes (PEUK), and potential mixotrophs (MEUK) (Fig. 2). Chlorophyll-containing eukaryotes were further subdivided into 2 size classes, pico (0.8-2 µm) and nano (>2-20 µm). HEUK were characterized by presence of LTG fluorescence and very low chlorophyll fluorescence (i.e., similar to surface PRO). PEUK had chlorophyll but little LTG fluorescence. MEUK had chlorophyll and LTG fluorescence.

**Figure 2.**
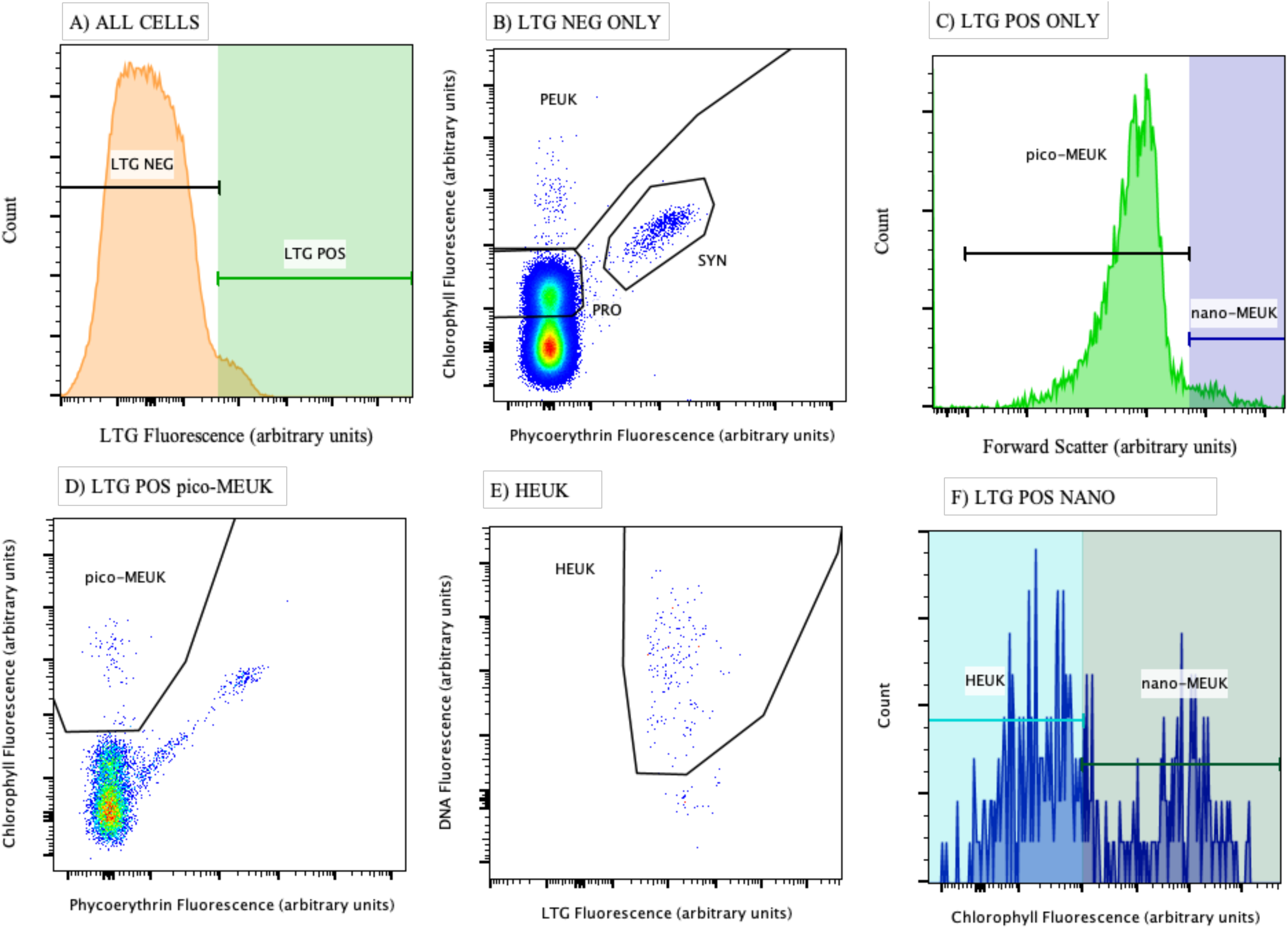
Flow cytometric gating scheme for delineating various eukaryotic populations. A) All cells are divided into LTG positive (LTG POS) and LTG negative (LTG NEG) using the maximum negative signal of co-occurring *Prochlorococcus* and heterotrophic bacteria, B) LTG NEG chlorophyll-bearing eukaryotic cells further refined by removing any prokaryotic populations using a Chl*a* vs. phycoerythrin plot, C) LTG POS cells divided into 2 size classes (pico- and nano-MEUK) using the forward light scatter signal of 1 µm polystyrene beads, D) LTG POS picoeukaryotes (pico-MEUK) further refined to remove any remaining prokaryotic cells using a Chl*a* vs. phycoerythrin plot, E) Heterotrophic Eukaryote (HEUK) cells shown as DNA vs. LTG fluorescence, F) LTG POS nanoeukaryotes divided into those with more chlorophyll fluorescence than *Prochlorococcus* (nano-MEUK) and those with less (HEUK).

The flow cytometric gating scheme was as follows: LTG positive and LTG negative particles were separated assuming that the maximum negative signal was that of co-occurring prokaryotes (PRO and HBACT). Thus, all particles with LTG signals greater than these prokaryotes were designated LTG positive (Fig. 2A). LTG negative particles were separated into prokaryotes and eukaryotes by examining their signatures on a chlorophyll vs. phycoerythrin plot, with eukaryotic cells having more chlorophyll than PRO or SYN (Fig. 2B). Eukaryotic cells without acidic vacuoles were assumed to be PEUK.

LTG positive and LTG negative particles were further subdivided into rough size categories relative to the forward light scatter (FSC) signal of a 1-µm fluorescent polystyrene bead (Polysciences, Inc.). Particles with FSC less than the 1-µm bead were designated pico-sized cells (Fig. 2C). Those with higher scatter were designated nano-sized cells (Fig. 2C). Due to their high refractive index, polystyrene fluorescent beads scatter more light than biological cells of similar size, such that the FSC dividing criterion for 1-µm beads is closer in size to a 2 or 3-µm cell (Spinrad and Brown, 1986; Chandler et al., 2011).

Due to fluorescence overlap of the LTG signal with phycoerythrin, a high percentage of the SYN usually registered as LTG positive even though they have no digestive vacuoles. Similarly, a small percentage (2-5%) of PRO and HBACT had to be gated out of the LTG positive category due to fluorescence overlap. Therefore, a further gating step was used to separate any remaining prokaryotes from LTG positive eukaryotic cells (Fig. 2D).

Once all prokaryotes were omitted from pico-sized LTG positive cells, the remaining cells were designated as potential mixotrophic picoeukaryotic cells (pico-MEUK) (Fig. 2D). Nano-sized LTG-positive cells were plotted as a single parameter histogram of chlorophyll fluorescence, and those with more chlorophyll than PRO were considered potential mixotrophic nano-eukaryotic cells (nano-MEUK) (Fig. 2F), whereas nano-eukaryotic LTG positive particles with very low chlorophyll were designated as HEUK (Fig. 2E, F).

To better compare LTG fluorescence between analysis days (when new dye batches were prepared), we normalized the LTG fluorescence of LTG positive groups to the mean LTG fluorescence of LTG negative groups run the same day. Specifically, LTG fluorescence of MEUK populations were normalized by dividing by the mean LTG fluorescence of PEUK, and LTG fluorescence of HEUK was normalized by dividing by the mean LTG fluorescence of HBACT.

### 2.3 Epifluorescence microscopy analyses

Water samples (450 mL) for epifluorescence microscopy were preserved with sequential additions of alkaline Lugol’s solution, borate-buffered formalin, and sodium thiosulfate, then stained with proflavine (Sherr and Sherr, 1993; Taylor et al., 2015). One subsample (50 mL) was prepared for nano-plankton analysis by filtering through a 0.8-µm pore-size black PCTE filter, staining with 4’, 6-diamidino-2 phenylindole (DAPI) prior filtering to dryness and mounting on a glass slide between immersion oil and a coverslip. The rest of the preserved samples (400 mL) was treated similarly but filtered through an 8-µm pore-size black PCTE filter for micro-plankton analysis. Slides were then frozen (-80°C) for shore-based imaging.

An Olympus Microscope DP72 Camera mounted on an Olympus BX51 fluorescence microscope was used to image each slide, using 60X magnification for the 0.8-µm filters (2-12 µm cells) and 20X magnification for the 8-µm filters (>12 µm cells) to capture 20 images per slide. Images were processed using ImageJ image analysis software (v 1.53c) to estimate abundance and biovolume (further details in Yingling et al., this issue).

### 2.4 Microplankton Biomass Estimates

Flow cytometry and microscopy data were used to estimate carbon biomass for each group. PRO and SYN abundances were converted to carbon as 32 and 101 fg C cell^-1^, respectively (Garrison et al., 2000). Picoeukaryotes, <2-µm MEUK and PEUK, were estimated to be 320 fg C cell^-1^ based on the mean biovolume (BV) for 0.8-1.9 µm diameter cells converted to carbon using Menden-Deuer and Lessard’s (2000) equation (i.e., pg C cell^-1^ = 0.216 x BV^0.939^). Nanoeukaryote carbon estimates for chlorophyll-bearing cells are also from the Menden-Deuer and Lessard (2000) equation applied to the mean biovolumes of 2-10 µm cells measured by epifluoresence microscopy (Yingling et al., this issue). This resulted in mean carbon estimates of 11 pg C cell^-1^ (Cycle 1), 9 pg C cell^-1^ (Cycle 2 and 4) and 5 pg C cell^-1^ (Cycle 3) for the nano-sized cells that contained chlorophyll. Epifluorescence microscopy also showed that mean HEUK (2-10 µm) equivalent spherical diameter was 3.74 µm, with little difference between cycles, which resulted in using 5 pg C cell^-1^ using the Menden-Deuer and Lessard (2000) biovolume-biomass equation.

## 3. Results

### 3.1 Study Area Characteristics

Station locations are listed in Table 1, along with dates of occupation, mixed layer depths (MLD, defined as the depth where density is 0.01 kg m^-3^ higher than the mean of the upper 5 m), and mean mixed-layer temperature, nitrate and phosphate concentrations. Overall, mixed layer waters were warm (28.5-30.6°C) and nutrient poor, with dissolved nitrate levels mostly below the detection limit of 0.01 to 0.03 µM and dissolved phosphate from <0.01 to 0.12 µM (Table 1). Density and fluorescence traces (CTD sensor data) are shown in Fig. 3, as well as MLD and euphotic zone depths (to 1% incident irradiance). All stations had pronounced deep chlorophyll maxima.

**Table 1.** Station characteristics where live samples were collected for LysoTracker Green staining. All CTD casts were conducted in February, 2022 at ∼18:00-19:00 UTC (02:00-03:00 local time) at the location shown. Shown are Cycle number, Date (Month-Day), Cast (CTD) number, Latitude (Lat, °N), Longitude (Long, °E), Mixed Layer Depth (MLD, m) calculated as 0.01 kg m^-3^ higher than mean of upper 5 m; Temperature (T, °C) equal to mean of MLD; and mean nitrate (NO_3_, µM) and phosphate (PO_4_, µM) in mixed layer, “-“ = no data; BDL = below detection limit of 0.01 µM.

| CYCLE | Date | CAST | LAT | LONG | MLD | T (°C) | NO<sub>3</sub> | PO<sub>4</sub> |
| --- | --- | --- | --- | --- | --- | --- | --- | --- |
| 1 | 02-04 | 18 | -15.3720 | 114.3663 | 42 | 28.57 | 0.01 | 0.09 |
|  | 02-05 | 25 | -15.3968 | 114.2331 | 54 | 28.53 | BDL | 0.11 |
|  | 02-06 | 33 | -15.4683 | 114.1920 | 30 | 28.57 | 0.01 | 0.12 |
|  | 02-07 | 38 | -15.5984 | 114.1351 | 26 | 28.62 | 0.03 | 0.06 |
| 2 | 02-12 | 68 | -16.9995 | 116.0730 | 11 | 28.84 | 0.01 | 0.05 |
| 3 | 02-15 | 86 | -15.9428 | 115.8287 | 8 | 29.44 | BDL | 0.04 |
|  | 02-16 | 93 | -15.8807 | 115.8381 | 6 | 29.91 | 0.01 | 0.03 |
|  | 02-17 | 99 | -15.7641 | 115.7990 | 6 | 30.05 | 0.01 | 0.05 |
|  | 02-18 | 106 | -15.6003 | 115.7757 | 7 | 30.18 | - | - |
| 4 | 02-20 | 114 | -15.8853 | 118.1423 | 12 | 30.50 | 0.02 | 0.05 |
|  | 02-21 | 121 | -15.9089 | 118.0893 | 12 | 30.53 | 0.01 | 0.11 |
|  | 02-22 | 128 | -15.9392 | 118.1204 | 16 | 30.55 | 0.03 | BDL |

**Figure 3.**
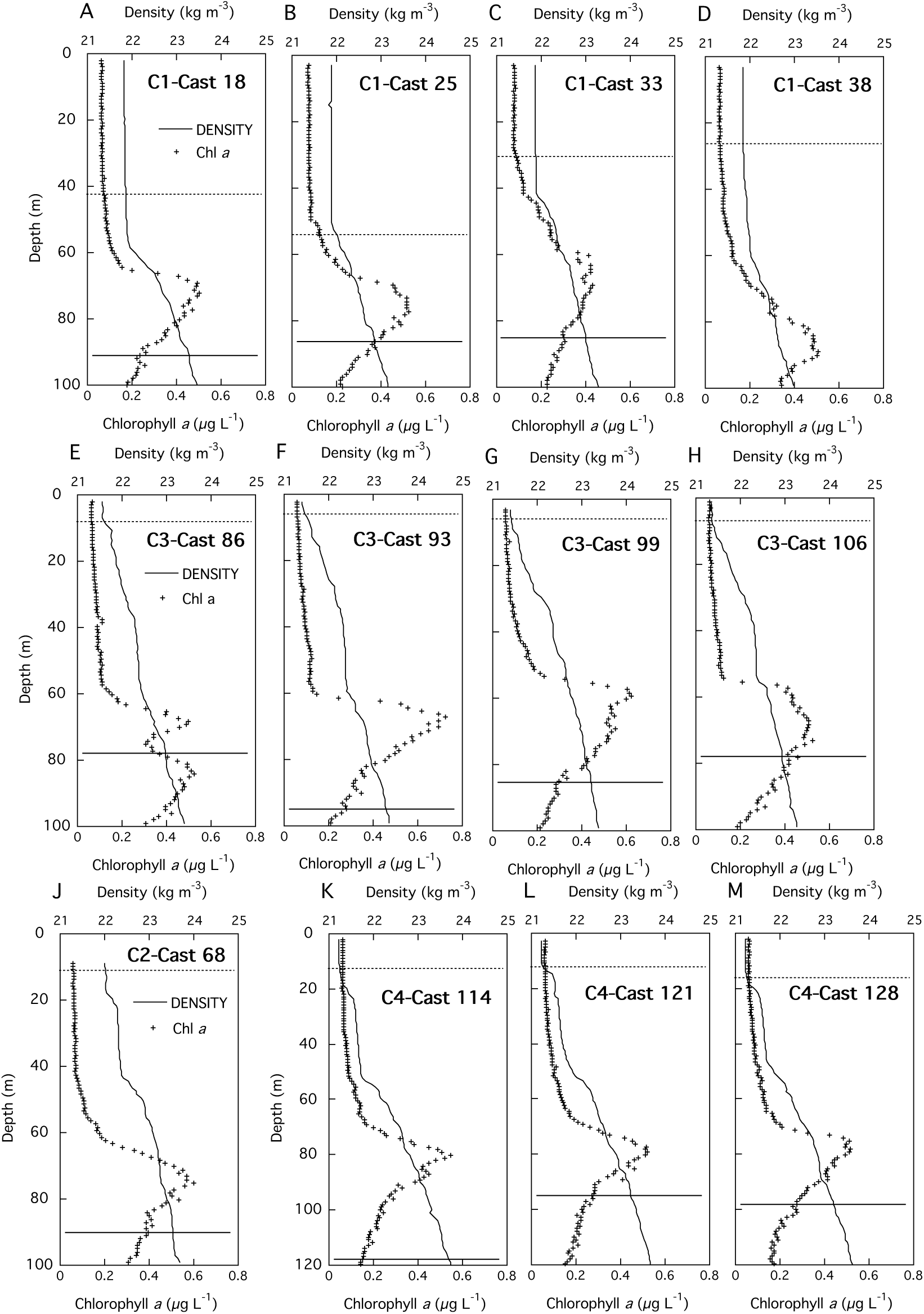
Density (kg m^-3^) and chlorophyll a (Chl*a*, µg L^-1^) fluorescence as a function of depth (m) for each cast in all Cycles. Also shown are the mixed layer depth (dotted line) and euphotic zone depth (to 1% incident irradiance, solid line), except for Cast 38 where no data for the exact euphotic zone was gathered before leaving the area (however likely ∼95 m from fluorescence trace). Cycle 1 (C1) data are shown in panels A-D; Cycle 3 (C3) in panels E-H, Cycle 2 (C2) in panel J, and Cycle 4 (C4) in panels K-M. Note that C4 depth axes are to 120 m, whereas all others are to 100 m.

Cycle 1 stations, occupied from 3-7 Feb 2022, experienced storm-induced mixing just before our occupation, resulting in changing mixed layer during the cycle (28-54 m). Euphotic zones extended to at least 85 m, below the deep chlorophyll maxima (65-90 m, Fig. 3A-D). CTD sensor traces of these properties showed very low fluorescence in the mixed layer (∼0.05 µg Chl*a* L^-1^), increasing to ∼0.4-0.5 µg Chl*a* L^-1^ at the deep chlorophyll maxima. Cycle 3 was occupied one week after Cycle 1, in the place that a marker drogue had drifted to from the last Cycle 1 station (Table 1). Cycle 3 stations were much more stratified, with shallower (6-8 m) and warmer (29-30°C) mixed layers. Cast 86, the first station of Cycle 3, had a double deep chlorophyll maximum, presumably formed from tidal action or local mixing of 2 water masses (Fig. 3E). Other stations in Cycle 3 had deep chlorophyll maxima between 60-70 m (Fig. 3F-H). Cycles 2 and 4 were also quite stratified, with very shallow mixed layers (≤16 m) and warm mean surface temperatures (29-30°C).

### 3.2. Eukaryotic abundances

Eukaryote distributions in each cycle are subdivided into different populations based on their presumed trophic state, and plotted as a function of depth (Fig. 4). Overall, HEUK averaged (± 1 s.e.) 524 ± 36 cells mL^-1^, when all depths and cycles were averaged. HEUK changed little with depth, and ranged from ∼300-600 cells mL^-1^ in surface samples, with a mean in the upper 40 m of 537 ± 43 cells mL^-1^. However, HEUK abundances in Cycles 1 and 3 had subsurface maxima near the deep chlorophyll maxima, while Cycle 2 showed no increase in abundance at that depth and Cycle 4 had a subsurface maximum shallower (∼55 m) than the deep chlorophyll maximum (∼80 m).

**Figure 4.**
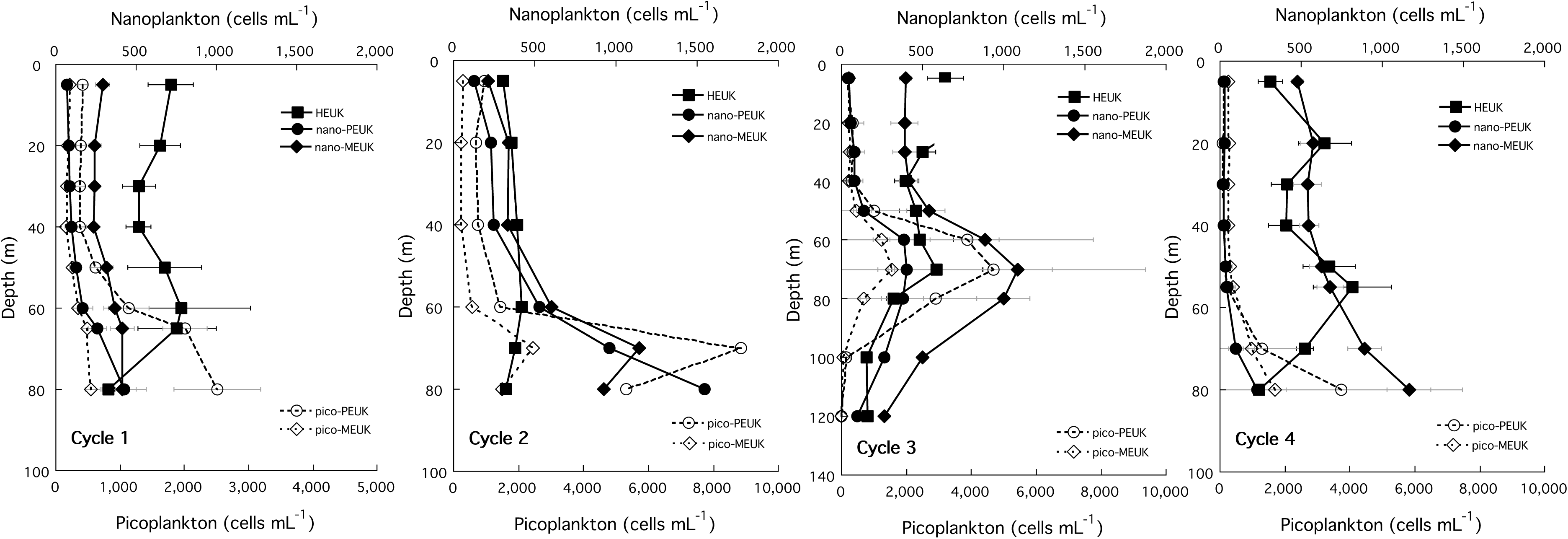
Eukaryotic plankton (cells mL^-1^, average ± 1 standard error) as a function of depth in each cycle. Populations shown are picoplankton (pico-PEUK and pico-MEUK) and nanoplankton (HEUK, nano-PEUK, and nano-MEUK). Note change of picoplankton x-axis in left-most figure.

In every cycle, nano-MEUK had higher abundances than nano-PEUK, with the exception of one sample in Cycle 2 at 80 m (Fig. 4; n.b., only one profile is available for Cycle 2 as no other casts had live samples analyzed). In the upper 40 m of Cycles 1, 3, and 4, nano-PEUK averaged 63 ± 5 cells mL^-1^, with a higher mean abundance in Cycle 2 (203 ± 38 cells mL^-1^). In contrast, nano-MEUK averaged 383 ± 22 cells mL^-1^ in these 3 cycles, with Cycle 2 having somewhat fewer nano-MEUK at 294 ± 41 cells mL^-1^. At depths greater than 40 m, nano-MEUK and nano-PEUK increased in abundance to the deep chlorophyll maximum, with means of 702 ± 56 and 324 ± 54 cells mL^-1^, respectively (Fig. 4).

In the upper 40 m, pico-PEUK were ∼4.5 times more abundant than nano-PEUK (325 ± 37 vs. 72 ± 9 cells mL^-1^, respectively), but nano-MEUK were more numerous than pico-MEUK (374 ± 24 vs. 233 ± 8 cells mL^-1^, respectively, Fig. 4). Like the nanoplankton, both pico-PEUK and pico-MEUK increased with depth, with means of 2544 ± 366 and 855 ± 103 cells mL^-1^, respectively, at depths >40 m to the deep chlorophyll maximum.

Partitioning the chlorophyll-bearing cells into presumed trophic status and size classes, reveals that the PEUK percentage as a function of depth largely mimics the shape of the Chl*a* fluorescence trace, with a maximum at depth (Fig. 5). On average, PEUK were 59 ± 0.03% (Cycle 1), 67 ± 0.02% (Cycle 2), 47 ± 0.05% (Cycle 3), and 25 ± 06% (Cycle 4) of the community. Considering Cycles 1, 3, and 4, MEUK were a higher proportion of the community in shallower waters, with the most variable portion being nano-MEUK. In Cycle 2, MEUK were a smaller proportion of the community (33 ± 0.02%) and that proportion varied little with depth.

**Figure 5.**
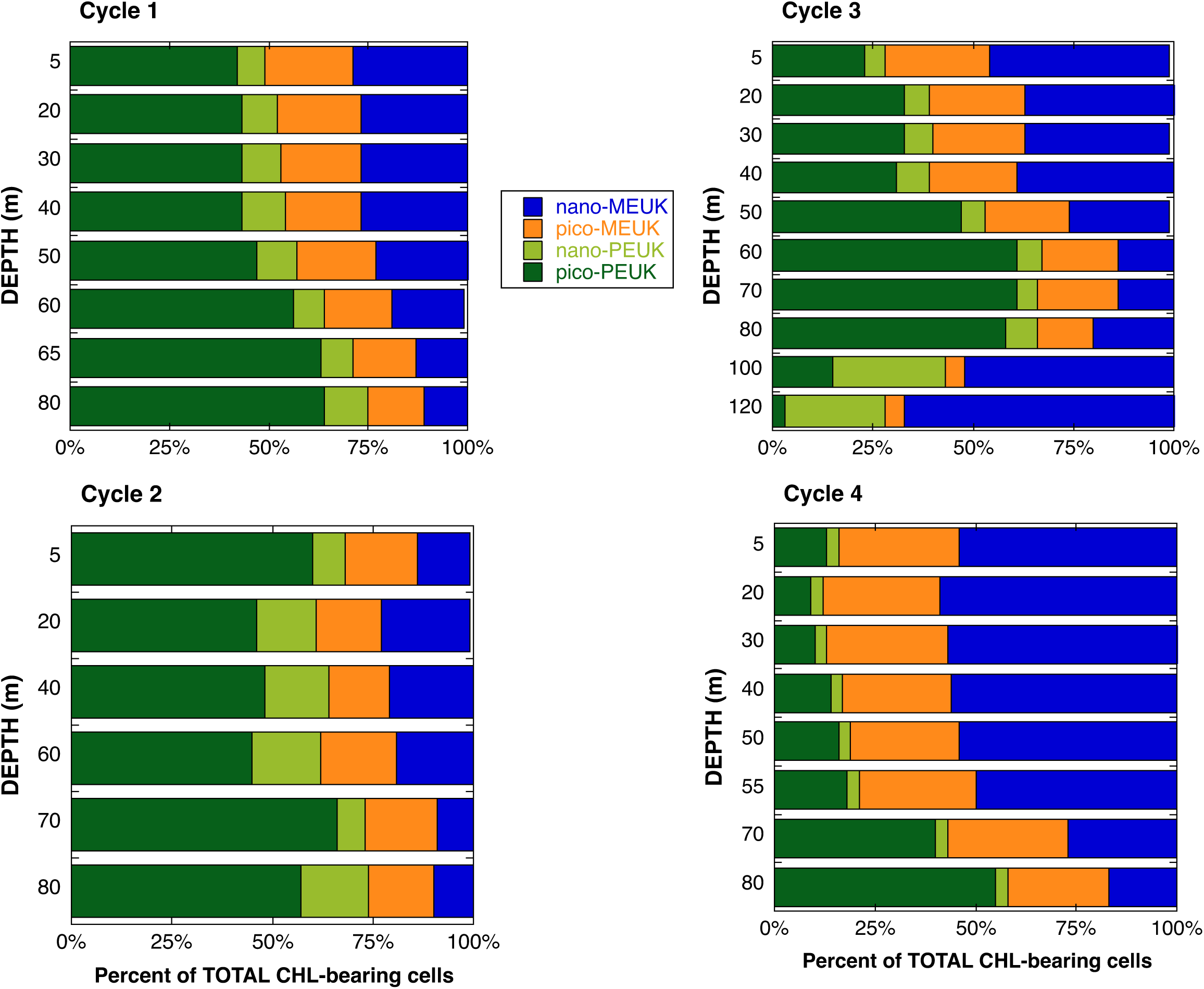
Proportion (%) of chlorophyll-bearing cell types at each depth (m) in each cycle. Shown are the percent of the total of pico and nano-MEUK, as well as pico- and nano-PEUK.

### 3.3 Microbial Biomass

All populations, including prokaryotes enumerated in the same samples, were converted to carbon biomass and integrated from 0-80 m (Table 2). HBACT represented ∼16% of the total microbial biomass, ranging from 14-17% in all cycles. PRO biomass accounted for 58% of the autotrophic community overall, with the lowest biomass contribution in Cycle 2 (38%) and the highest in Cycle 1 (57%). Nano-MEUK were the second highest contributor to autotrophic biomass, at 28% overall, followed by nano-PEUK at 9%, with pico-PEUK, pico-MEUK and SYN equaling only 6% of the total autotrophic community biomass. HEUKs were ∼13% of total microbial biomass, with a high of 17% in Cycle 1 and a low of 9% in Cycle 2. Overall, acid vacuole containing cells (MEUK plus HEUK) were 80% of the eukaryotic microbial biomass, ranging from 58% in Cycle 2 to 93% in Cycle 4. Details of autotrophic microbial biomass are discussed by Selph et al. (this issue, b)

**Table 2.** Carbon biomass in each cycle (integrals, 0-80 m, mg C m^-2^) of all populations enumerated with flow cytometry on live (unpreserved), Hoechst (DNA) and LysoTracker Green (acidic vacuole) stained cells. Also shows are the mean ± 1 standard error (s.e.) of each cycle’s data. Populations shown are *Prochlorococcus* (PRO), *Synechococcus* (SYN), heterotrophic (non-pigmented) bacteria (HBACT), obligate pico (pico-PEUK) and obligate nano (nano-PEUK) phytoplankton, as well as mixotrophic pico (pico-MEUK) and nano (nano-MEUK) phytoplankton, and heterotrophic eukaryotes (HEUK).

| CYCLE | CAST | PRO | SYN | HBACT | pico-PEUK | nano-PEUK | pico-MEUK | nano-MEUK | HEUK |
| --- | --- | --- | --- | --- | --- | --- | --- | --- | --- |
| **1** | 18 | 580 | 13 | 215 | 25 | 150 | 11 | 396 | 169 |
|  | 25 | 638 | 12 | 221 | 24 | 133 | 7 | 254 | 190 |
|  | 33 | 511 | 9 | 198 | 22 | 154 | 5 | 247 | 285 |
|  | 38 | 719 | 14 | 216 | 24 | 98 | 7 | 212 | 391 |
| **Mean (1 s.e.)** |  | **612 (44)** | **12 (1)** | **213 (5)** | **24 (1)** | **134 (13)** | **8 (1)** | **277 (41)** | **259 (51)** |
| **2** | 68 | **484** | **28** | **239** | **56** | **323** | **17** | **363** | **144** |
| **3** | 86 | 863 | 32 | 333 | 36 | 43 | 15 | 603 | 303 |
|  | 93 | 798 | 25 | 314 | 51 | 108 | 16 | 343 | 156 |
|  | 99 | 685 | 35 | 298 | 41 | 86 | 12 | 313 | 194 |
|  | 106 | 693 | 21 | 305 | 23 | 34 | 13 | 517 | 182 |
| **Mean (1 s.e.)** |  | **759 (43)** | **28 (3)** | **312 (8)** | **38 (6)** | **68 (17)** | **14 (1)** | **444 (70)** | **209 (32)** |
| **4** | 114 | 710 | 26 | 294 | 16 | 34 | 16 | 575 | 267 |
|  | 121 | 651 | 18 | 265 | 16 | 52 | 11 | 443 | 191 |
|  | 128 | 627 | 20 | 280 | 10 | 31 | 11 | 402 | 159 |
| **Mean (1 s.e.)** |  | **663 (24)** | **21 (2)** | **280 (8)** | **14 (2)** | **39 (7)** | **13 (1)** | **473 (52)** | **206 (22)** |
| **OVERALL Mean (1 s.e.)** |  | **663 (31)** | **21 (2)** | **265 (13)** | **29 (4)** | **104 (24)** | **12 (1)** | **315 (36)** | **219 (22)** |

### 3.4 Flow Cytometry cell parameters

Comparing LTG fluorescence between eukaryotic groups in terms of arbitrary fluorescence units (f.u.) per cell (Table 3) showed a large increase over the signals of organisms that do not possess acidic vacuoles (i.e., PRO and HBACT). Pico- and nano-PEUK had low LTG fluorescence (587 and 989 f.u. cell^-1^, respectively). MEUK had much higher LTG fluorescence, at 27,000 f.u. per pico-MEUK and 280,000 f.u. nano-MEUK cell. HEUK also had high mean LTG fluorescence (24,000 f.u. cell^-1^), though not as high as the larger nano-MEUK. Fluorescence values substantially above our set threshold levels are viewed as simply indicating the presence of digestive vacuoles and phagotrophic functionality and do not necessarily relate to digestive efficiency or capacity.

**Table 3.** LysoTracker Green (LTG) fluorescence per cell in each microbial group as measured by flow cytometry. The units are fluorescence per cell, and are arbitrary, as this parameter is not calibrated to any standard. Microbial groups are: *Prochlorococcus* (PRO), heterotrophic bacteria (HBACT), obligate picophytoplankton (pico-PEUK), obligate nanophytoplankton (nano-PEUK), mixotrophic picoplankton (pico-MEUK), mixotrophic nanoplankton (nano-MEUK), and heterotrophic eukaryotes (HEUK). Shown are the values as averages ± 1 standard error (N=76), pooling all data from every cycle and depth.

| Microbial Group | LTG Fluorescence cell<sup>-1</sup> |
| --- | --- |
| PRO* | 570 ± 95 |
| HBACT | 217 ± 29 |
| Pico-PEUK | 587 ± 53 |
| Nano-PEUK | 989 ± 59 |
| Pico-MEUK | 27,551 ± 2,325 |
| Nano-MEUK | 282,848 ± 33,386 |
| HEUK | 24,338 ± 1,074 |

\**Prochlorococcus* maximum value: 3349 f.u. cell<sup>-1</sup>

The mean LTG fluorescence in the HEUK and MEUK groups were divided by the group’s mean chlorophyll fluorescence (LTG:Chl*a*) and plotted as a function of depth (Fig. 6). This ratio was over 40-times lower for pico- and nano-PEUK (0.004 ± 0.004 and 0.003 ± 0.001, mean ± 1 s.e., respectively), so these groups are not shown. No consistent depth pattern was found for the LTG:Chl*a* ratio for the HEUK, but in Cycles 1-3, the ratio averaged 7.5 ± 0.4, with a 2.7-11.1 range. In Cycle 4, the LTG:Chl*a* ratio was higher for HEUKs (10.1 ± 1.6; range 5.1-33.2). Pico-MEUK usually had a higher ratio of LTG:Chl*a* near the surface, decreasing with depth, driven by photoadaption as cellular chlorophyll increased with decreasing light. Overall, the average LTG:Chl*a* ratio for pico- and nano-MEUK was 0.17 ± 0.03 and 0.22 ± 0.03, respectively, which is ∼40-50-times lower than for HEUKs.

**Figure 6.**
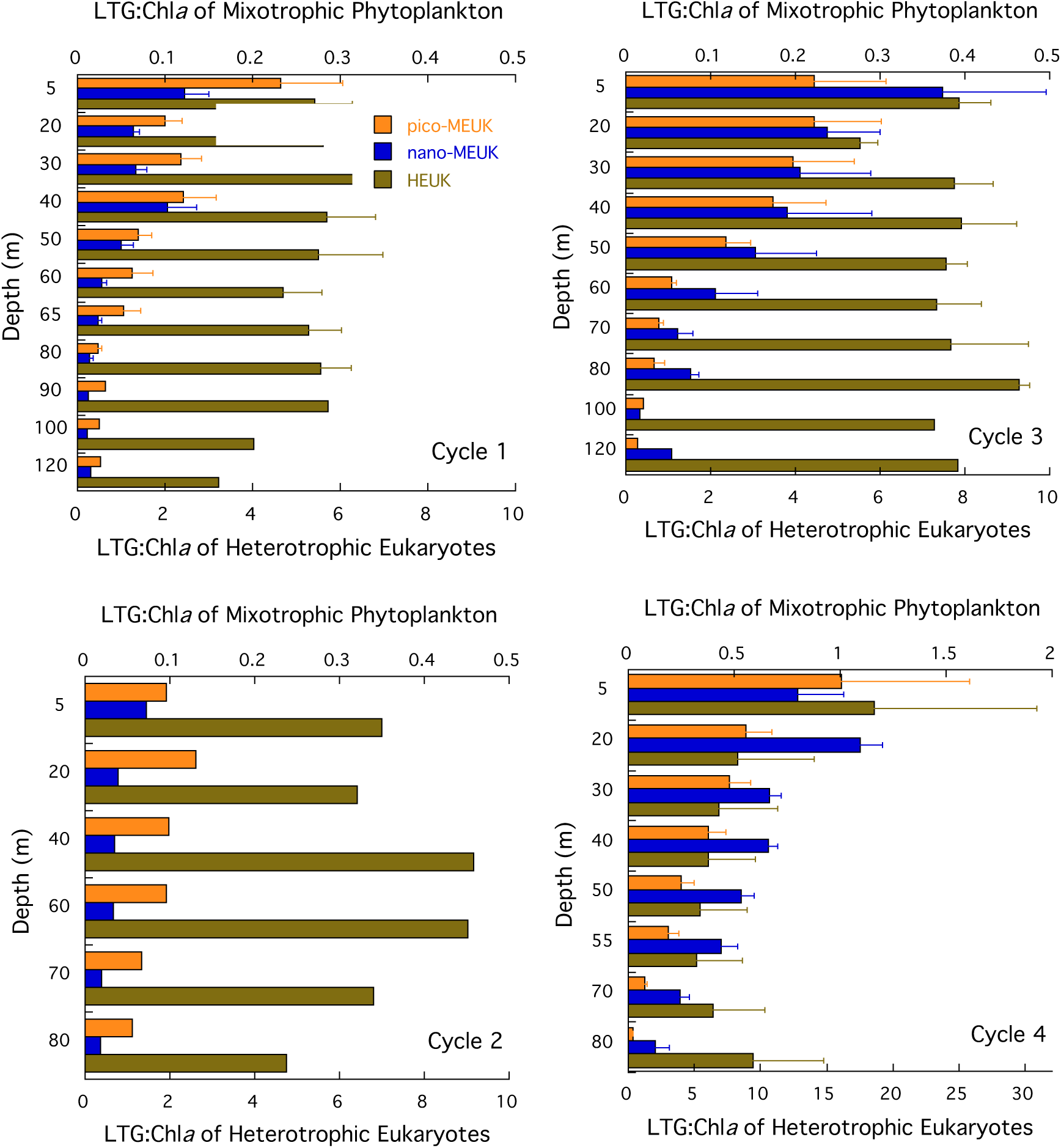
Mean LTG fluorescence:chlorophyll fluorescence in acid-vacuole containing groups as a function of depth for each cycle. Shown are pico- and nano-MEUK, as well as HEUK.

The mean LTG fluorescence relative to FSC (proxy for cell size) was also plotted as a function of depth (Fig. 7). As expected, the LTG:FSC is close to zero (0.02 ± 0.002) for PEUK, since they have no acidic vacuoles. The HEUK LTG:FSC ratio averages 0.13 ± 0.01 for all depths and cycles. Pico-MEUK and nano-MEUK averaged 1.08 ± 0.08 and 1.11 ± 0.12 overall, however in the Cycle 4 averages for these groups was higher than the other cycles (1.60 ± 0.29 and 2.67 ± 0.21, respectively). Also, LTG:FSC tended to be higher at the surface than near the DCM for the MEUK groups (Fig. 7).

**Figure 7.**
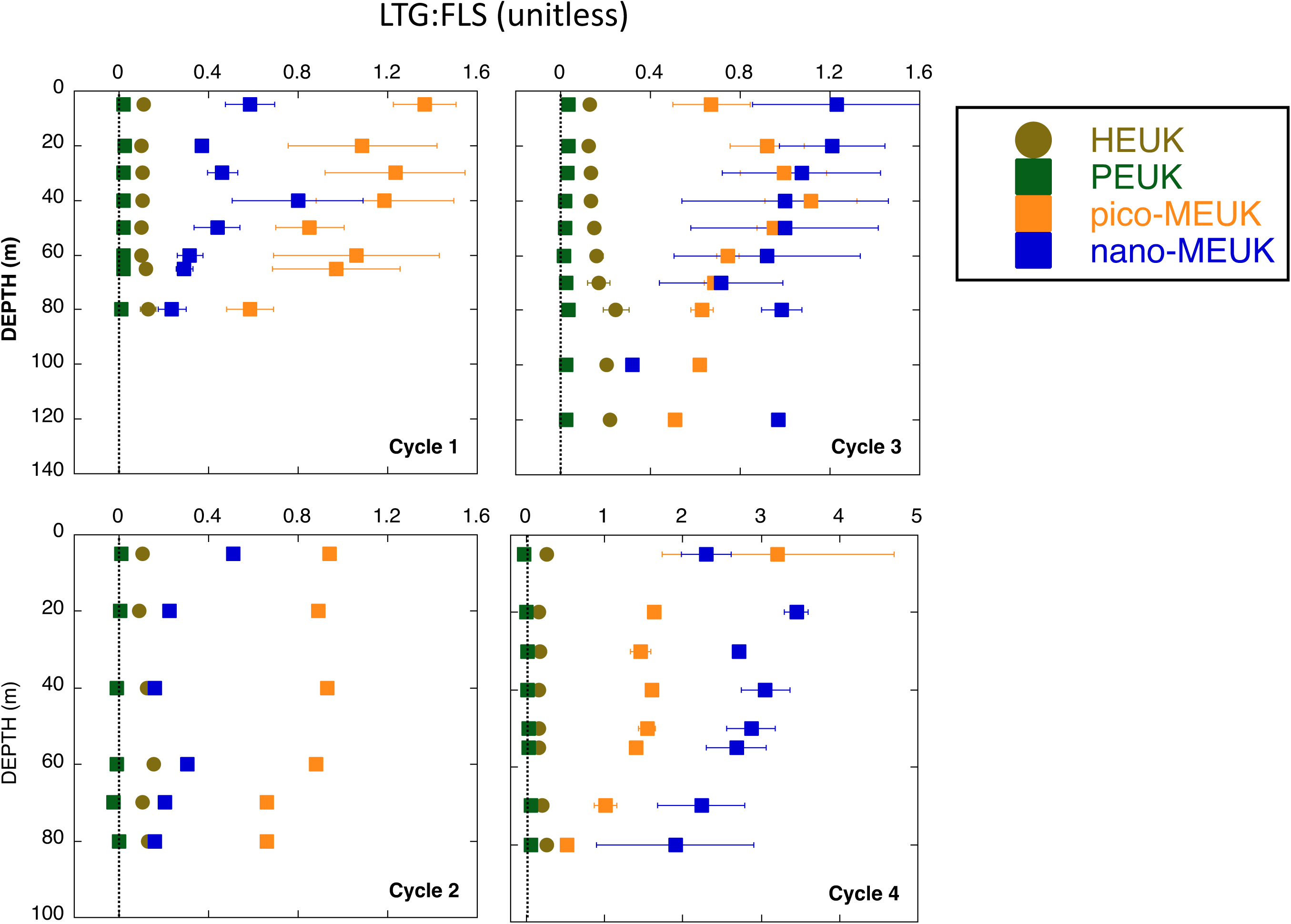
Mean LTG fluorescence:forward light scatter in all eukaryotic groups, as a function of depth for each cycle. Shown are HEUK, PEUK, pico- and nano-MEUK.

## 4. Discussion

We sampled the microbial community of an open-ocean oligotrophic region using shipboard flow cytometry with the LysoTracker Green stain to evaluate phagotrophic function among chlorophyll-containing (mixotrophs) and non-chlorophyll-containing (heterotrophs) cells. A high proportion of the chlorophyll-containing cells were found to be mixotrophs. In the following subsections, we consider potential methodological issues in this analysis, mixotroph abundance and biomass contributions the microbial community, inferred predator-prey relationships, and comparisons to other data sets.

### 4.1 Methodological Concerns

Multi-step gating was used to separate cells into functional categories, based on a combination of scatter, chlorophyll, phycoerythrin, LTG and DNA fluorescence. Previous reports (e.g., Rose et al., 2004; Sintes and del Giorgio, 2010) established that LTG does not accumulate appreciably in prokaryotic cells, and our work confirmed this. We therefore used the LTG (green) fluorescence signal from PRO and non-pigmented bacteria as the threshold level for LTG POS cells.

Rose et al. (2004) indicated that LTG stained all acidic vacuoles, including chloroplasts of cultured protists, *Micromonas pusilla* and *Heterocapsa triquetra*, that were being grown phototrophically. Others have reported no appreciable staining of small obligate autotrophs or cells grown under autotrophy-ideal conditions (Anderson et al., 2017; Costa et al., 2022). In Rose et al. (2004), the presumed chloroplast staining of *Heterocapsa triquetra* could have captured some vacuole activity, even though this functionally mixotrophic dinoflagellate was not actively feeding at the time. Our data suggest that some chloroplast staining by LTG may occur, as the magnitude of LTG fluorescence for plastidic cells, especially nano-MEUK, was much higher than for the aplastidic HEUK. If this is so, a varying threshold may be needed to account for chloroplast and increasing cell size effects.

Of additional concern for non-specific staining, Wilken et al. (2019) found LTG staining in the frustules of diatoms, and Sato and Hashihama (2019) noted that high LTG staining specifically occurred at stations where small (<10 µm) pennate diatoms were abundant. Thus, in natural samples with significant diatom abundance, interpretation of LTG-stained chlorophyll-containing cells would have to account for this effect. In our case, we found little evidence of diatoms from either microscopy (Yingling et al., this issue) or HPLC pigment data (Selph et al., this issue, b). It is therefore unlikely that our data reflect misidentification of diatoms as mixotrophs.

Another caveat about these results is that some fraction of the cells may be undergoing apoptosis (cell death) and degrading their own chloroplasts in acidic vacuoles (Koester et al., 2024). Thus, the absolute abundance of mixotrophs from LTG staining must be considered as an upper bound. Assessment of apoptosis in parallel with LTG-staining might allow estimation of this process in natural samples.

We found background levels of LTG staining (∼40-times lower signal) for the photoeukaryotic group that we designated as obligate phototrophs (PEUKs). Among LTG-positive cells, an unknown fraction of cells designated MEUK could be HEUK with partially digested autotrophic prey in them. The majority, however, must be true mixotrophs given their high chlorophyll fluorescence and the fact that acidic vacuoles are processed quickly, such that little LTG signal would be expected in cells that are not actively phagocytizing prey. The same issues also apply to distinguishing groups by fluorescence microscopy unless vacuoles are large enough to view chlorophyll-bearing prey within them. Additionally, our samples were taken at night (∼02:30 local, analyzed before dawn), potentially a time of more active feeding by mixotrophs as opposed to photosynthesis during daytime. There is contradictory evidence on whether heterotrophic eukaryotes feed more during the day than night (Strom, 2001; Arias et al., 2020;), as well as some evidence of nocturnal feeding by mixotrophs (Porter, 1988; Anderson et al., 2017). Future studies might examine LTG staining of natural protistan communities over the full photoperiod to assess diel patterns and possible differences between heterotrophic and mixotrophic components.

Another potential methodological concern is the incubation time of LTG with a sample prior to analyzing on the flow cytometer. Rose et al. (2004) recommended a short incubation time (10 min) to detect maximal concentrations of lysotracker-stained vacuoles for a ciliate and flagellate at culture concentrations of 10^4^ to 10^5^ cells mL^-1^, which were 10 to 100 times denser than our natural samples (∼10^3^ cells mL^-1^). Given our much lower densities and therefore longer analysis times to count an appropriate number of cells (all were analyzed within 1 h after staining), we could not follow the Rose at al. (2004) protocol precisely. However, we did a test for time sensitivity of LTG fluorescence brightness and found no appreciable difference between the same samples run after 10 versus 60 min of staining (data not shown).

Estimating cell size with flow cytometry using polystyrene beads can also be somewhat problematic. Rose et al. (2004) used a 2.5 µm bead as a FSC cutoff for delineating eukaryotes from prokaryotes, with particles larger than that included as eukaryotes and smaller ones excluded. There are 2 issues with using bead size as a cutoff. First, because light scatter in flow cytometry is not an exact indicator of size and because solid fluorescent beads have higher light scatter signals (due to their higher refractive index) than same-sized biological cells (Wang and Hoffman, 2017), the 2.5-µm size cutoff would likely exclude 2.5-µm cells (and any smaller eukaryotes), and only represent much larger cells. Second, many different taxa of picoeukaryotic flagellates occur in open ocean waters, many of which are approximately 2-3 µm in size. In choosing a 1-µm bead to distinguish between pico and nanoeukaryotes, we assumed that cells scattering more light than a 1-µm bead were the full eukaryote component (after eliminating PRO and SYN in the same samples using chlorophyll and phycoerythrin signals).

We dual stained these samples with LTG and a DNA stain (Hoechst 34580), and used the DNA stain to separate the HBACT, PRO and SYN from the eukaryotes. To our knowledge, this is the first report of dual staining the same sample to get both signals from individual cells. Unlike other green nuclear stains, such as SybrGreen, staining with LTG did not affect the intrinsic chlorophyll or phycoerythrin fluorescence of cells, however the green fluorescence from LTG did overlap with the orange fluorescence of phycoerythrin from SYN, so SYN had to be gated out (in separate chlorophyll vs. phycoerythrin plots) so as not to include those cells as LTG positive plankton.

### 4.2 Mixotroph contribution to total microbial populations

Acid vacuole-containing organisms comprised the majority of eukaryotic biomass (mixotrophs plus heterotrophs) and about 1/3 of total microbial biomass (prokaryotes + eukaryotes). Stated another way, presumptive mixotrophs alone were about 1/3 of total autotrophs, with PRO having the most biomass of any autotrophic group.

In support of our findings on mixotrophy, the dominant groups of chlorophyll-bearing eukaryotes from high performance liquid chromatography (HPLC) were prymnesiophytes, dinoflagellates, prasinophytes, pelagophytes and cryptophytes, all of which have mixotrophic members (Selph et al., this issue, b). Corroborating these pigment results, we found that the dominant genera from analyses of relative abundances of 18S rRNA genes were also mostly those with mixotrophic species (Selph et al., this issue, b). DNA data showed dominance of cryptophytes, known mixotrophs, and the dominant dinoflagellate genera all have mixotrophic species (*Gyrodinium*, *Gymnodinium, Prorocentrum, Warnowia, Karlodinium, Lepidodinium, Tripos* and *Margalefidinium*). Mixotrophic species were also found in the dominant prasinophyte genera detected - *Ostreococcus and Bathycoccus*. Other mixotroph dominants were *Parmales,* and the prymnesiophyte genus *Chrysochromulina*. Conversely, high relative abundances of taxa generally not known to have mixotrophic species were the prymnesiophyte *Phaeocystis* and the pelagophyte *Pelagomonas caleolata*. Note, however, that one species of *Phaeocystis* has been observed with ingested bacteria by Koppelle et al. (2022), and pelagophytes were more prevalent near the base of the euphotic zone where fewer mixotrophs were present in our study.

Pico-MEUK and nano-PEUK were both relatively stable portions of the community, with the nano-MEUK showing the most change with depth in the water column. In addition, the proportion of MEUKs relative to all PEUK as a function of forward light scatter of chlorophyll-bearing cells shows a strong positive correlation (r^2^ = 0.83, Fig. 8). This suggests that larger autotrophs are more likely to be mixotrophic than smaller autotrophs, a result also found by Sato and Hashihama (2019) for North Pacific Ocean/Bering Sea populations. Thus, abundance changes in nano-MEUK with depth might reflect their responses to proximate nutrient or light conditions. Along with this, cell size as trait has been theorized to influence whether a cell is better fit for mixotrophy vs. obligate phototrophy, as smaller cells (picoplankton) get higher fluxes of light and nutrients (higher surface:volume ratio), whereas larger cells can acquire more nutrients from phagotrophy than diffusion (Ward and Follows, 2016; Millette et al., 2023).

**Figure 8.**
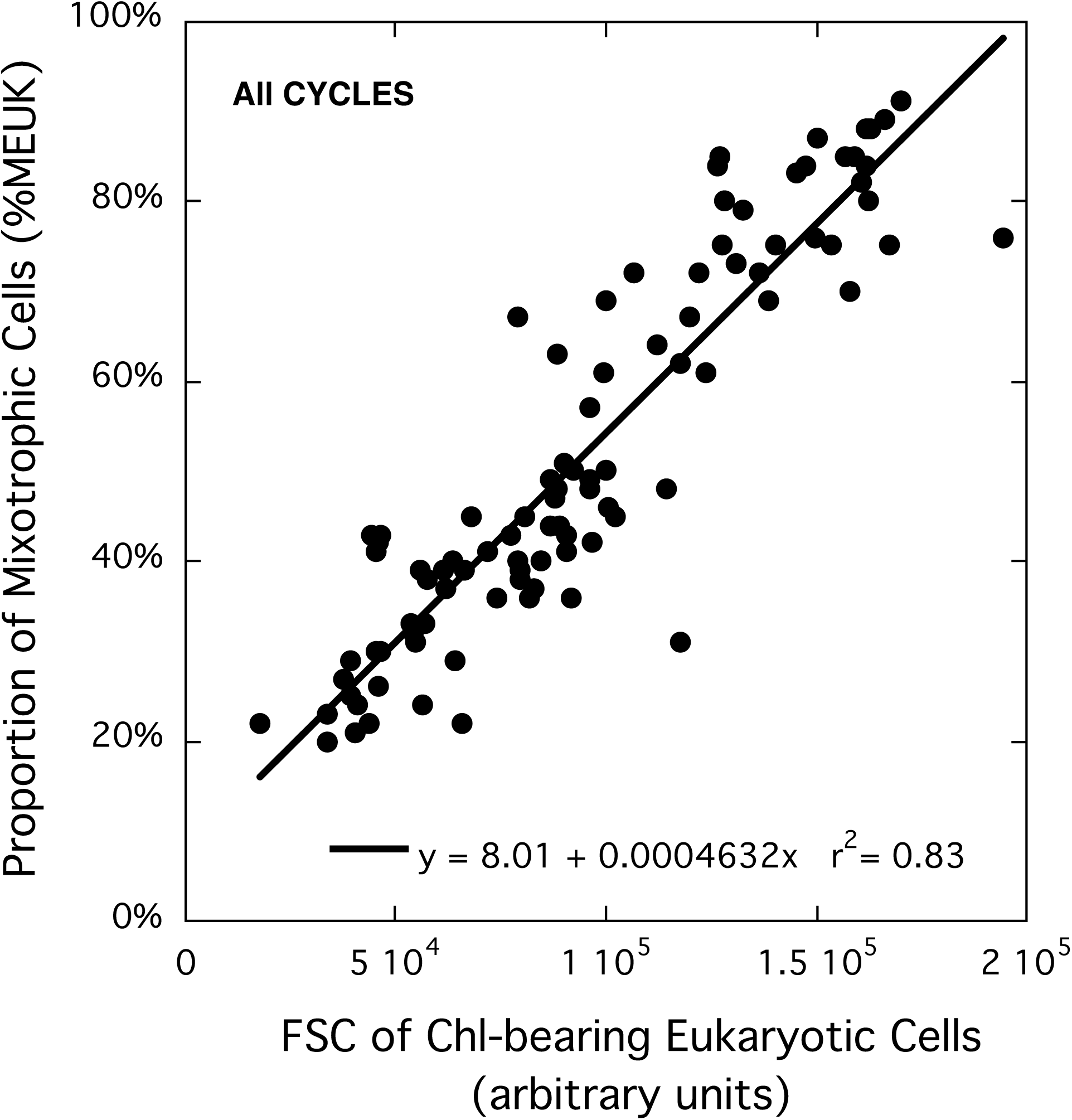
Proportion of mixotrophic cells (%MEUK) as a function of the forward light scatter of chlorophyll-bearing cells.

Mixotrophs were a higher proportion of the autotrophic community at shallower depths with high light and low nutrients, consistent with a nutrient-acquisition argument for their prevalence. A comparable study by Sato and Hashihama (2019) found that the percent of mixotrophs increased in phosphorus-poor areas, similar to our results (although our limiting nutrient is likely nitrogen). Other research has suggested that a combination of high light and low nutrients favor mixotrophy as a strategy (Hartmann et al., 2012). Along with this, pico- and nano-PEUKs were a majority of the chlorophyll-bearing cells at the deep chlorophyll maxima – a depth that coincides with the nutricline, and thus represents the depth where nutrient limitation is likely overtaken by light limitation as the driving factor for cell growth. In this study, the deep chlorophyll maxima were also biomass maxima for chlorophyll-containing eukaryotes, not just pigment maxima.

Metabolic theory suggests that mixotrophs should become more phagotrophic as temperature increases (Wilken et al., 2013). Consistent with this, over the 23.2 to 30.6 °C temperature range of our euphotic zone samples, the proportion of chlorophyll-containing eukaryotes classified as mixotrophic increased with as higher temperature (r^2^ = 0.636, P<0.001). However, this correlation likely just reflects the fact that the warmer temperatures represented the shallower depths with high light levels and lowest nutrients that select for mixotrophy.

### 4.3 Grazer and prey inferences

Our study area was dominated by picoplanktonic organisms, notably PRO and HBACT. SYN and picoeukaryotes combined represented <10% of PRO biomass. Thus, as confirmed by epifluorescence microscopy, most of the microbial grazers (HEUK and nano-MEUK) seen by flow cytometry would likely be nanoflagellates to handle such small prey. We postulate that HEUK principally consumed PRO, SYN or HBACT, since the amount of Chl*a* they had was equal or less than that of PRO cells. Some of the cells classified as a mixotrophs could, however, be heterotrophic cells containing larger plastidic eukaryote prey, multiple ingested PRO or ingested SYN, which we found to register as LTG positive due to their phycoerythrin pigment.

While correlation is not causation, we found that HEUK biomass had no significant relationship (at the 0.01 level) with the biomass of any group except for PRO (r^2^ = 0.368, P<0.01), suggesting PRO was a significant component of their diet, as would be intuitively the case based on numerical abundance of PRO. The lack of correlation with HBACT is somewhat surprising, especially given reviews such as Sanders et al. (1992), which showed positive correlations between these parameters from many studies. However, it may be that any such relationship is obscured in our data set, since PRO dominated microbial biomass (41% vs. 16% for PRO vs. HBACT, respectively).

In contrast, we found no significant correlations between mixotrophs and PRO biomass. This does not preclude some grazing of mixotrophs on PRO, as Hartmann et al. (2013) found, and laboratory studies by Li et al. (2020) demonstrated. Nonetheless, it suggests that mixotroph grazing on PRO is not likely a significant mortality agent for PRO during our study. However, significant positive correlations between MEUK and HBACT were found (pico-MEUK: r^2^ = 0.613, p<0.001; nano-MEUK: r^2^ = 0.638, p<0.001). This is consistent with other studies which found significant bacterivory by mixotrophs (Havskum and Riemann, 1996; Hartmann et al., 2012; Sanders and Gast, 2012; Unrein et al., 2014), which is postulated to be an acquisition strategy for limiting nutrient, particularly under high light conditions. For instance, Zubkov and Tarran (2008) showed that <5-µm phototrophs were responsible for a high proportion of the bacterivory in the nitrogen-limited euphotic zone of the North Atlantic Ocean and the tropical North-East Atlantic Ocean, although their impact on standing stock was low. Similarly, high bacterivory by small phototrophic flagellates in the northwest Mediterranean Sea might have been a P-acquisition strategy (Unrein et al., 2007).

### 4.4 Comparison to other data sets

A global biogeographic analysis revealed that constitutive mixotrophs smaller than 20-µm were under-sampled and under-represented in ocean data bases (Leles et al., 2018). Therefore, studies like this one that distinguish acid vacuole-containing organisms can help address this knowledge gap. Published reports using LTG to identify mixotrophs are most commonly from surface samples in coastal or river environments, and thus not directly comparable to our study (e.g., Rose et al., 2004; Sintes and Del Giorgio, 2010; Heywood et al., 2011; Anderson et al., 2017). However, Sato and Hashihama (2019) used LTG on depth-resolved samples in the North Pacific Ocean/Bering Sea, reporting on latitudinal transects at 10 m and June-Aug 2014 and at five euphotic zone depths for Aug-Oct 2017. Presumptive mixotrophs ranged from 20-70% of pico- and nanophytoplankton abundance, with a higher percentage in the subtropical gyre (the closest conditions to our study) and lower in equatorial and subarctic regions (except for the Bering Sea). Livanou et al. (2019) results compared well to our upper 40 m means of 537 and 383 cells mL^-1^ for HEUK and nano-MEUK, respectively, finding approximately 500 and 350 cells mL^-1^ for <5 µm HEUKs and nano-MEUK, respectively, at 5 m in oligotrophic waters of the Mediterranean Sea.

Using fluorescently-labelled bacteria (FLB) as prey surrogates has shown mixotrophs to be important consumers of picoplankton. In nutrient-depleted surface waters of the Bay of Aarhus (Denmark), for example, Havskum and Riemann (1996) found that ∼half of the pigmented nanoflagellates were mixotrophs, whereas <10% were seen below the pycnocline, suggesting that low nutrients encouraged this trophic mode. Sanders et al. (2000) found 7-18% of chlorophyll-bearing nanoflagellates to be mixotrophs in the upper 10-20 m of the Sargasso Sea. Unrein et al. (2007) reported that mixotrophs contributed ∼50% to total flagellate grazing at an oligotrophic coastal station in the Mediterranean Sea, and Unrein et al. (2014) found haptophytes to be the most important bacterivorous mixotrophs over the annual cycle. In contrast, Anderson et al. (2017) followed FLB-uptake seasonally in surface samples of a harbor in Denmark, finding a large range of mixotroph concentration (5,000-40,000 cells mL^-1^) but low overall percentage (15%) of actively feeding cells. Similarly, Christaki et al. (1999) examined depth profiles of nanoflagellates and found little bacterivory by mixotrophs in the oligotrophic Aegean Sea (5% of bacterial production) relative to heterotrophic nanoflagellates (40%); however, the labeled prey may have been too large for most of the nanoflagellates to handle. We note that the mixotroph concentration in Christaki et al. (1999) study were 5 to 10 times lower than heterotrophic nanoflagellates, unlike more similar abundances in the present study. Based on the results of such studies, the high proportion of acid vacuole-containing cells and presumptive mixotrophs in our samples suggests a significant grazing role for them in the oligotrophic Argo Basin.

### 4.5 Summary and Conclusions

This study used shipboard flow cytometry to characterize phagotrophic functionality within the microbial community in open-ocean waters of Argo Basin off NW Australia. We stained unpreserved (live) samples with an acid lysozyme stain (LysoTracker Green) and a DNA stain (Hoechst 34580), and we enumerated heterotrophic bacteria, heterotrophic eukaryotes, and the chlorophyll-bearing cells of PRO, SYN, pico- and nano-eukaryotes. Pico- and nano-eukaryotes were further separated into obligate phototrophs and mixotrophs by the absence or presence of acidic vacuoles. Waters of the Argo Basin are warm, stratified and oligotrophic, with PRO playing a major role in primary production. The eukaryotic phytoplankton were comprised of roughly 40% (abundance) or 70% (biomass) of mixotrophs, which were significantly correlated with heterotrophic bacteria, their presumed main prey. Heterotrophic eukaryotes, mainly nanoflagellates, showed little variation in abundance with depth, and were significantly correlated with PRO. The combination of mixotrophs and heterotrophs, both acid vacuole-containing groups, represented the majority of the eukaryotic microbial community. These results highlight the key role of grazing in an oceanic oligotrophic region, with most protists augmenting their nutrition by phagotrophy.

## Acknowledgements

We thank the captain and crew of the R/V *Roger Revelle* for their excellent ship-based support, use of scientific collection equipment, and assistance with logistics. Research support was provided by the National Science Foundation Grants OCE-1851381, OCE-1851347 and OCE-1851558. Seawater and plankton samples were collected under Australian Government permit AU-COM2021-520 and Australian Marine Parks permit PA2021-00062-2 issued by the Director of National Parks, Australia. The views expressed in this publication do not necessarily represent those of the Director of National Parks or the Australian Government.

## Author statement

K.E.S. developed the protocols for mixotrophy measurements by flow cytometry, conducted field sampling, analyzed results and drafted the manuscript. N.Y., C.T. and M.R.L. conducted field sampling and provided feedback on concepts in the manuscript. All authors contributed to comments and edits of the manuscript.

## References

Allen, R.D., Fok, A.K., 1983. Nonlysosomal vesicles (acidosomes) are involved in phagosome acidification in *Paramecium*. J. Cell Biol. 97, 566–570.

Anderson, R., Jurgens, K., Hansen, P.J., 2017. Mixotrophic phytoflagellate bacterivory field measurements strongly biased by standard approaches: A case study. Frontiers in Microbiol. 8, doi: 10.3389/fmicrb.2017.01398.

Becker, S., Aoyama, M., Woodward, E.M.S., Bakker, K., Coverly, S., Mahaffey, C., Tanhua, T., 2019. The precise and accurate determination of dissolved inorganic nutrients in seawater; Continuous Flow Analysis methods and laboratory practices, IOCCP Report No. 14, GO-SHIP Repeat Hydrography Manual: A Collection of Expert Reports and Guidelines, ICPO Publication Series No. 134.

Chandler, W.L, Yeung, W., Tait, J.F., 2011. A new microparticle size calibration standard for use in measuring smaller microparticles using a new flow cytometer. J. Thromb. Haemostasis 9(6), 1216–1224.

Christaki, U., Van Wambeke, F., Dolan, J.R., 1999. Nanoflagellates (mixotrophs, heterotrophs and autotrophs) in the oligotrophic eastern Mediterranean: standing stocks, bacterivory and relationships with bacterial production. Mar. Ecol. Prog. Ser. 181, 297–307.

Costa, M.R.A., Sarmento, H., Becker, V., Bagatini, I.L., Unrein, F., 2022. Phytoplankton phagotrophy across nutrients and light gradients using different measurement techniques. J. Plank. Res. 44(4), 508–521. doi: 10.109/plankt/fbac035.

Flynn, K.J., Mitra, A., Anestis, K., Anschütz, A.A., Calbet, A., Ferreira, G.D., Gypens, N., Hansen, P.J., John, U., Martin, J.L., Mansour, J.S., 2019. Mixotrophic protists and a new paradigm for marine ecology: where does plankton research go now? J. Plank. Res. 41(4), 375–91.

Fok, A.K., Lee, Y., Allen, R.D., 1982. The correlation of digestive vacuole pH and size with the digestive cycle in *Paramecium caudatum*. J. Protozool. 29(3), 409–414.

Garrison, D.L., Gowing, M.M., Hughes, M.P., Campbell, L., Caron, D.A., Dennett, M.R., Shalapyonok, A., Olson, R.J., Landry, M.R., Brown, S.L., Liu, H.-B., Azam, F., Steward, G.F., Ducklow, H.W., Smith, D.C., 2000. Microbial food web structure in the Arabian Sea: a US JGOFS study. Deep-Sea Res. II 47, 1387–1422.

Hartmann, M., Grob, C., Tarran, G.A., Martin, A.P., Burkill, P.H., Scanlan, D.J., Zubkov, M.V., 2012. Mixotrophic basis of Atlantic oligotrophic ecosystems. Proc. Natl. Acad. Sci. 109, 5756–5760.

Hartmann, M., Zubkov, M.V., Scanlan, D.J., Lepere, C., 2013. In situ interactions between photosynthetic picoeukaryotes and bacterioplankton in the Atlantic Ocean: evidence for mixotrophy. Environ. Microbiol. Rep. 5, 835–840.

Havskum, H., Riemann, B., 1996. Ecological importance of bacterivorous, pigmented flagellates (mixotrophs) in the Bay of Aarhus, Denmark. Mar. Ecol. Prog. Ser. 137,251–263.

Heywood, J.L., Sieracki, M.E., Bellows, W., Poulton, N.J., Stepanauskas, R., 2011. Capturing diversity of marine heterotrophic protists: one cell at a time. ISME J. 5, 674–684.

Koester, J.A., Fox, O., Smith, E., Cox, M.B., Taylor, A.R., 2024. A multifunctional organelle coordinates phagocytosis and chlorophagy in a marine eukaryote phytoplankton Scyphosphaera apsteinii. New Phytol. 246, 1096–1112.

Koppelle, S., López-Escardó, D., Brussaard, C.P., Huisman, J., Philippart, C.J., Massana, R., Wilken, S., 2022. Mixotrophy in the bloom-forming genus *Phaeocystis* and other haptophytes. Harmful Algae 117, 102292.

Landry, M.R., Ohman, M.D., Goericke, R., Stukel, M.R. and Tsyrklevich, K., 2009. Lagrangian studies of phytoplankton growth and grazing relationships in a coastal upwelling ecosystem off Southern California. Prog. Oceanogr. 83(1-4), 208–216.

Landry, M.R., Stukel, M.R., Yingling, N., Selph, K.E., Kranz, S.A., Fender, C.K., Swalethorp, R., Bhabu, R.I., This issue. Microbial food web dynamics in tropical waters of the bluefin tuna spawning region off northwestern Australia. Deep-Sea Res. II.

Lefort, T., Gasol, J.M., 2014. Short-time scale coupling of picoplankton community structure and single-cell heterotrophic activity in winter in coastal NW Mediterranean Sea waters. J. Plank. Res. 36, 243–258.

Legendre, L., Courties, C., Troussellier, M., 2001. Flow cytometry in oceanography 1989–1999: Environmental challenges and research trends. Cytometry, 44(3), 164–172.

Leles, S.G., Mitra, A., Flynn, K.J., Tillmann, U., Stoecker, D., Jeong, H.J., Burkholder, J., Hansen, P.J., Caron, D.A. Glibert, P.M., Hallegraeff, G., Raven, J.A., Sanders, R.W., Zubkov, M., 2019. Sampling bias misrepresents the biogeographical significance of constitutive mixotrophs across global oceans. Global Ecol Biogeogr. 28, 418–428.

Li, Q., Edwards, K.F., Schvarcz, C.R., Selph, K.E., Steward, G.F., 2021. Plasticity in the grazing ecophysiology of *Florenciella* (Dichtyochophyceae), a mixotrophic nanoflagellate that consumes *Prochlorococcus* and other bacteria. Limnol. Oceanogr. 66(1), 47–60.

Livanou, E., Lagaria, A., Santi, I., Mandalakis, M. Pavlidou, A. Lika, K. Psarra, S., 2019. Pigmented and heterotrophic nanoflagellates: abundance and grazing on prokaryotic picoplankton in the ultra-oligotrophic eastern Mediterranean Sea. Deep-Sea Res. II 164, 100– 111, doi:10.1016/j.dsr2.2019.04.007.

Menden-Deuer, S., Lessard, E.J., 2000. Carbon to volume relationships for dinoflagellates, diatoms, and other protist plankton. Limnol. Oceanogr. 45, 569–679.

Millette, N.C., Gast, R.J., Luo, J.Y., Moeller, H.V., Stamieszkin, K., Andersen, K.H., Brownlee, E.F., Cohen, N.R., Duhamel, S., Dutkiewicz, S., Glibert, P.M., 2023. Mixoplankton and mixotrophy: future research priorities. J. Plank. Res. 45(4), 576–96.

Mills, D.B., 2020. The origin of phagocytosis in Earth history. Interface Focus 10: 20200019, doi:10.109/rsfs.2020.0019.

Mitra, A., K.J. Flynn, J. M. Burkholder, T. Berge, A. Calbet, J.A. Raven, E. Graneli, P.M. Glibert, P. J. Hansen, D. K. Stoecker, F. Thingstad, U. Tillmann, S. Vage, S. Wilken, and M. V. Zukov, 2014. The role of mixotrophic protists in the biological carbon pump. Biogeosci. 11, 995–1005.

Rose, J.M, Caron, D.A., Sieracki, M.E., Poulton, N., 2004. Counting heterotrophic nanoplanktonic protists in cultures and aquatic communities by flow cytometry. Aquat. Microb. Ecol. 34, 263–277.

Sanders, R.W., Caron, D.A., Berninger, U.-G., 1992. Relationships between bacteria and heterotrophic nanoplankton in marine and fresh waters: an inter-ecosystem comparison. Mar. Ecol. Prog. Ser. 86, 1–14.

Sanders, R.W., Berninger, U.-G., Lim, E.L., Kemp, P.F., Caron, D.A., 2000. Heterotrophic and mixotrophic nanoplankton predation on picoplankton in the Sargasso Sea and on Georges Bank. Mar. Ecol. Prog. Ser. 192, 103–118.

Sanders, R.W., Gast, R.J., 2012. Bacterivory by phototrophic picoplankton and nanoplankton in Arctic waters. FEMS Microbiol. Ecol. 82, 242–253.

Sato, M., Hashihama, F., 2019. Assessment of potential phagotrophy by pico- and nanophytoplankton in the North Pacific Ocean using flow cytometry. Aquat. Microb. Ecol. 82, 275–288.

Selph, K.E., 2021. Enumeration of marine microbial organisms by flow cytometry using near- UV excitation of Hoechst 34580-stained DNA. Limnol. Oceanogr. Meth. 19, 692–701, 10.1002/lom3.10454.

Selph, K.E., Lampe, R.H., Yingling, N., Landry, M.R. This issue, b. Phytoplankton community composition and biomass in the oligotrophic Argo Basin (NW Australia). Deep-Sea Res. II.

Sherr, E.B., Sherr, B.F., 1993. Preservation and storage of samples for enumeration of heterotrophic protists. IN: Kemp, P.F., Sherr, B.F., Sherr, E.B., Cole, J.J. (Eds.), Current methods in aquatic microbial ecology. Lewis Pub., Boca Raton, FL, 207–212.

Sherr, E.B., Caron, D.A., Sherr, B.F., 2018. Staining of heterotrophic protists for visualization via epifluorescence microscopy. IN: Handbook of methods in aquatic microbial ecology, CRC Press, 213–227.

Sintes, E., del Giorgio, P.A., 2010. Community heterogeneity and single-cell digestive activity of estuarine heterotrophic nanoflagellates assessed using lysotracker and flow cytometry. Environ. Microbiol. 12(7), 1913–1925.

Spinrad, R.W., Brown, J.F., 1986. Relative real refractive index of marine microorganisms: a technique for flow cytometric estimation. Appl. Optics 25(12), 1930–1934.

Strom, S.L., 2001. Light-aided digestion, grazing and growth in herbivorous protists. Aquat. Microb. Ecol. 23, 253–261.

Taylor, A.G., Landry, M.R., Selph, K.E., Wokuluk, J.J., 2015. Temporal and spatial patterns of microbial community biomass and composition in the Southern California Current ecosystem. Deep Sea Res. II 112, 117–128.

Unrein, F., Massana, R., Alonso-Saez, L., Gasol, J.M., 2007. Significant year-round effect of small mixotrophic flagellates on bacterioplankton in an oligotrophic coastal system. Limnol. Oceanogr. 52, 456–469.

Unrein, F., Gasol, J.M., Not, F., Forn, I., Massana, R., 2014. Mixotrophic haptophytes are key bacterial grazers in oligotrophic coastal waters. ISME J. 8(1), 64–176.

Vazquez-Dominguez, E., Casamayor, E.O., Catala, P., Lebaron, P., 2005. Different marine heterotrophic nanoflagellates affect differentially the composition of enriched bacterial communities. Microb. Ecol. 49, 474–495.

Vives-Rego, J., Lebaron, P., Nebe-von Caron, G., 2000. Current and future applications of flow cytometry in aquatic microbiology, FEMS Microbiol. Rev., 24(4), 429–448, doi: 10.1111/j.1574-6976.2000.tb00549.x.

Wang, L., Hoffmann, R.A., 2017. Standardization, calibration, and control in flow cytometry. Curr. Protoc. Cytom. 79,1.3.1–1.3.27, doi: 10.1002/cpcy.14.

Ward, B.A., Follows, M.J., 2016. Marine mixotrophy increases trophic transfer efficiency, mean organism size, and vertical carbon flux. Proc. Natl. Acad. Sci. 113, 2958–2963.

Wilken, S., Huisman, J., Naus-Wiezer, S., Van Donk, E., 2013. Mixotrophic organisms become more heterotrophic with rising temperature. Ecol. Lett. 16, 225–233.

Wilken, S., Yung, C.C., Hamilton, M., Hoadley, K., Nzongo, J., Eckmann, C., Corrochano-Luque, M., Poirier, C., Worden, A.Z., 2019. The need to account for cell biology in characterizing predatory mixotrophs in aquatic environments. Phil. Trans. Roy. Soc. B, 374(1786), 20190090.

Yingling, N., Selph, K. E., Landry, M. R., Kranz, S. A., Johnson, M., Stukel, M. R., This issue. Phytoplankton nutrient uptake, abundance, biomass and community composition in the oligotrophic Argo Basin, Indian Ocean.

Zubkov, M.V., Tarran, G.A., 2008. High bacterivory by the smallest phytoplankton in the North Atlantic Ocean. Nature, 455, 224–227, doi:10.1038/nature07236.

